# Distinct neural signatures underlying information manipulation in working memory

**DOI:** 10.1101/2023.07.11.548635

**Authors:** Dongping Shi, Qing Yu

## Abstract

Working memory is a core cognitive ability that supports both the maintenance and manipulation of information in mind to serve complex behavior. Previously, stimulus representation in memory maintenance has been observed in both voltage and alpha-band activity in electroencephalography (EEG). However, the exact function of these two neural signatures in working memory has remained elusive. Here we systematically examined this question by asking whether memory manipulation is supported by the same neural signatures as memory maintenance. Human participants either maintained a previously-seen spatial location, or manipulated the spatial location according to a mental rotation cue over a delay. Using multivariate distance-based decoding, we observed robust neural representations of maintained and manipulated locations in both low-frequency voltage and alpha-band oscillatory activity, suggesting that memory maintenance and manipulation are supported by multiple overlapping neural signatures in EEG. Yet, these neural signatures demonstrated distinct spatiotemporal dynamics: location representations in alpha-band activity were most evident in the posterior channels, while location representations in voltage were most evident in the more anterior, central channels. The temporal emergence of manipulated representation in central voltage also preceded that in posterior alpha-band activity, suggesting that voltage might carry stimulus-specific source signals originated internally from more anterior cortex, whereas alpha-band activity might reflect feedback signals in posterior cortex received from higher-order cortex. Lastly, while location representations in both signals could be compressed into a low-dimensional neural subspace, location representation in central voltage was higher-dimensional and underwent a representational transformation that exclusively predicted memory behavior. Together, these results highlight the crucial role of central voltage in memory manipulation, as well as offer direct experimental evidence supporting functional distinction between voltage and alpha-band oscillatory activity in EEG.

## Introduction

Working memory, the ability to flexibly maintain and manipulate information in mind, is fundamental to high-level cognition, including decision making, reasoning, and arithmetic [1]. One central question regarding the neural mechanism of working memory is to understand the representation of stimulus information held in working memory. Over the past years, utilizing multivariate decoding or encoding techniques, converging evidence has revealed significant stimulus representations from delay-period activity in electroencephalography (EEG). Among these studies, two important neural signatures of memory maintenance have been identified: voltage and alpha-band (8-12 Hz) oscillatory activity. Both types of signals during memory delay represent stimulus information across a wide range of stimulus types, including spatial location [2,3], orientation [4,5], and even object [6–8].

However, the presence of stimulus information in both signals does not necessarily imply that these two signals serve identical function in working memory. In fact, recent work has suggested the functions of stimulus representation in voltage and alpha-band activity might significantly differ: for instance, one study showed that during memory maintenance, voltage reflected contents in working memory whereas alpha-band activity reflected spatial attention signals [4]. Moreover, two other studies showed that when attentional prioritization controlled the in and out of information from the focus of attention, only the attended information could be decoded from voltage [9] whereas the unattended information could be decoded from alpha-band activity [10]. In other words, while stimulus representations in voltage and alpha-band activity have greatly fructified our understanding of the neural mechanism of memory maintenance, the nature of these two signals in working memory remains to be explored.

On the other hand, an important hallmark of working memory is its capability of manipulating information following different behavioral goals. This cognitive flexibility of information manipulation distinguishes working memory from mere, passive maintenance of short-term memory. Critically, information derived from memory manipulation is internally generated, rather than externally perceived as in memory maintenance. This unique characteristic makes it an ideal tool to study the process of internal generation and maintenance of mnemonic information without contamination from sensory signals. However, although the neural mechanism of memory maintenance has been studied extensively, the neural mechanism underlying memory manipulation has not been well understood. A common approach to study information manipulation in working memory is to utilize a mental rotation paradigm. In this paradigm, participants first view a to-be-memorized item such as spatial location or orientation, and mentally rotate the item by a specific angle defined by a rotation cue. Using this paradigm, evidence from the functional magnetic resonance imaging (fMRI) literature has shown that the rotated information was decodable in early visual and parietal cortex, and shared neural codes with the maintained information [11,12]. More recent work using ultra-high field fMRI has further demonstrated that rotated information was predominantly represented in the superficial and deep layers of early visual cortex, suggesting the feedback nature of the rotated representations in visual cortex [13]. However, the low temporal resolution of fMRI has precluded further interpretation on this topic. Meanwhile, EEG offers excellent temporal resolution to address this question; however, research on memory manipulation using EEG has been rare. Two recent preprints [14,15] tested this question with EEG, by focusing on alpha-band and voltage activity, respectively. They found that although the rotated information was decodable in both alpha-band and voltage activity, it was generalizable to the maintained information in alpha-band but not in voltage activity. However, at this point, it remains to be further investigated whether this apparent discrepancy was due to functional differences between signals or differences between studies instead.

Here we sought to systematically examine this question by asking: 1) whether the two neural signatures of memory maintenance, namely voltage and alpha-band activity, also contributed to information representation during memory manipulation; and 2) whether and how stimulus representations in voltage and alpha-band activity differ between memory maintenance and manipulation; and 3) whether the spatiotemporal dynamics of rotated stimulus representations differ between voltage and alpha-band activity. Combining EEG, multivariate decoding, and state space analyses, we aimed to provide a thorough understanding of the functions of voltage and alpha-band activity, as well as their respective contributions to memory maintenance and manipulation.

## Results

### Behavioral performance

Participants (N = 49) performed a spatial working memory task while EEG was recorded (Figs 1A-B). The task consisted of two conditions: maintenance and manipulation. On each trial, participants first viewed one or two dots presented at different spatial locations, and were retrocued on the color of the to-be-maintained or manipulated dot. In the maintenance condition, participants were required to maintain the cued dot location throughout the delay. In the manipulation condition, participants were required to mentally rotate the cued dot location (either clockwisely or counterclockwisely) according to the cue during the delay.

**Fig 1.**
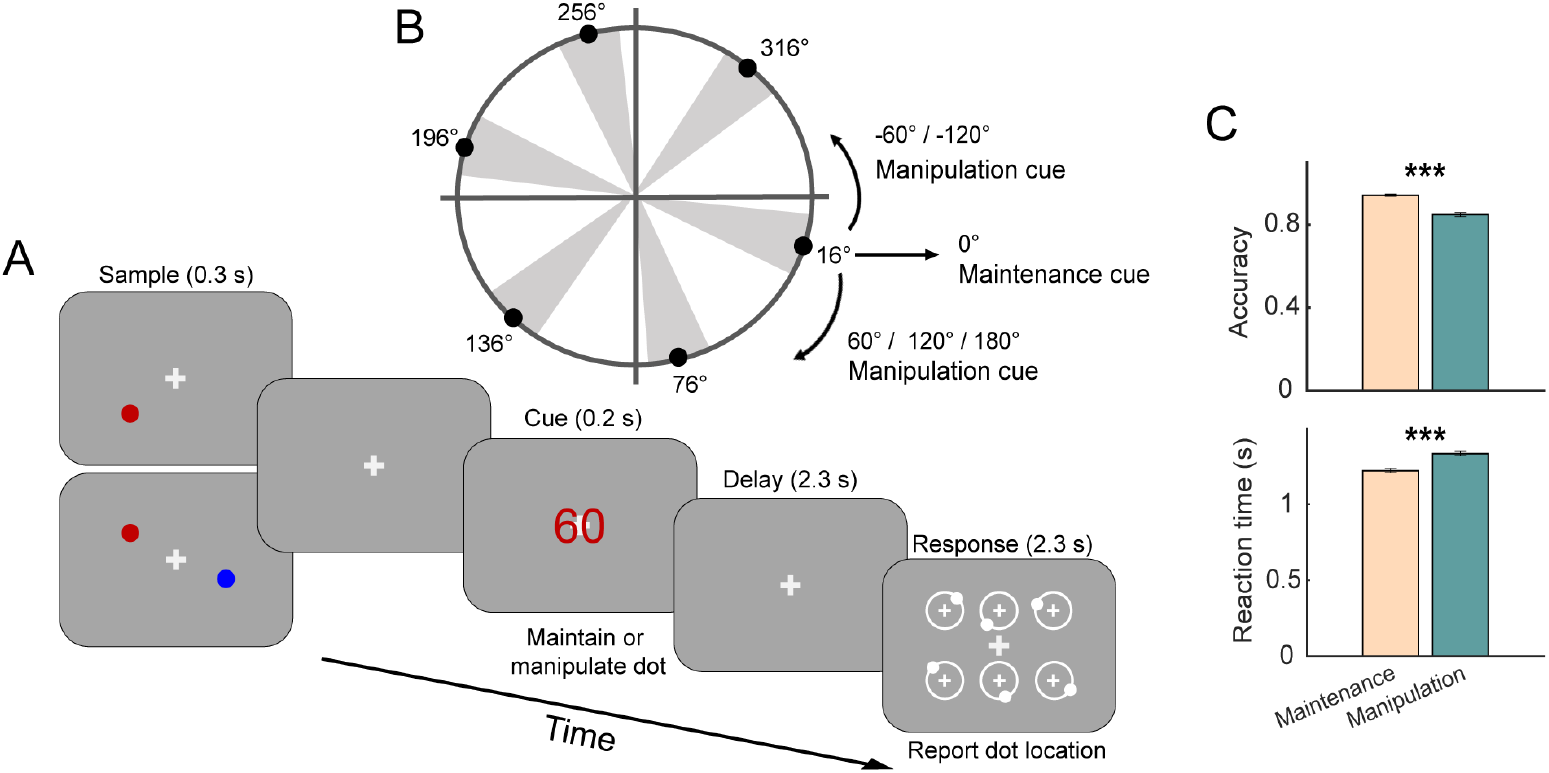
Behavioral paradigm and results. (A) Task schematics. The task consisted of two conditions: maintenance and manipulation. Participants first viewed either one or two dots presented at different spatial locations. A subsequent digit cue indicated the color of the cued dot, and whether participants needed to perform maintenance or manipulation on the cued dot. In the maintenance condition, the cue was always 0, and participants needed to maintain the dot location during the delay period. In the manipulation condition, the cue could be any one from ±60, ±120 or 180, and participants needed to manipulate the dot location with the cued color either clockwisely (positive values) or counterclockwisely (negative values) by the corresponding cued angle. During the response period, participants selected the correct dot location from six probes. (B) Distribution of sample dot locations. The dots’ locations were distributed across six bins, with a random jitter of ± 10° in each bin. (C) Accuracy and reaction time of behavioral performance in the maintenance and manipulation conditions. *** denotes *p* < 0.001. Error bars denote ±1 SEM.

Overall, participants performed both two conditions well, indicating their capability of performing fine-scaled mental rotation on spatial locations. In addition, performance in the maintenance condition was superior to that in the manipulation condition: the accuracy of behavioral performance in maintenance (*M* = 0.94, *SD* = 0.04) was significantly higher than that in manipulation (*M* = 0.85, *SD* = 0.07), *t* (48) = 13.68, *p* < 0.001. Likewise, reaction time in maintenance (*M* = 1.22, *SD* = 0.10) was also faster than that in manipulation (*M* = 1.33, *SD* = 0.11), *t* (48) = -17.36, *p* < 0.001 (Fig 1C).

### Spatiotemporal characterization of location representations in voltage and alpha-band activity during memory maintenance and manipulation

Previous studies have demonstrated reliable decoding of information held in working memory from both sustained potentials and alpha-band oscillatory activity in EEG [4,9,10,16]. Here we first tested whether the two neural signatures of memory maintenance also contributed to memory manipulation. We utilized a distance-based decoding approach [5,9] on all EEG electrodes to decode the to-be-maintained or manipulated location from either voltage or alpha-band activity across the trial time course. Each decoder was trained and tested on data within the same condition. All statistical significance for location decoding was evaluated by one-sample *t* tests and corrected for multiple comparisons using a cluster-based permutation method. Decoding of location displayed comparable time courses in voltage and alpha-band activity (Fig 2): in the maintenance condition, successful decoding of cued location emerged during the sample period, and sustained until the end of delay; In the manipulation condition, participants first viewed the cued location, and manipulated the cued location to the rotated location following the retrocue. As such, a tradeoff between decoding of cued and rotated locations was observed: decoding time course of the cued location was similar to that in maintenance during sample and early delay, but became progressively worse till the end of delay; by contrast, decoding of rotated location became increasingly evident during delay, along with the decrease in cued location decoding. These results suggested that location information in memory manipulation was represented in both voltage and alpha-band activity, in line with previous reports in memory maintenance.

**Fig 2.**
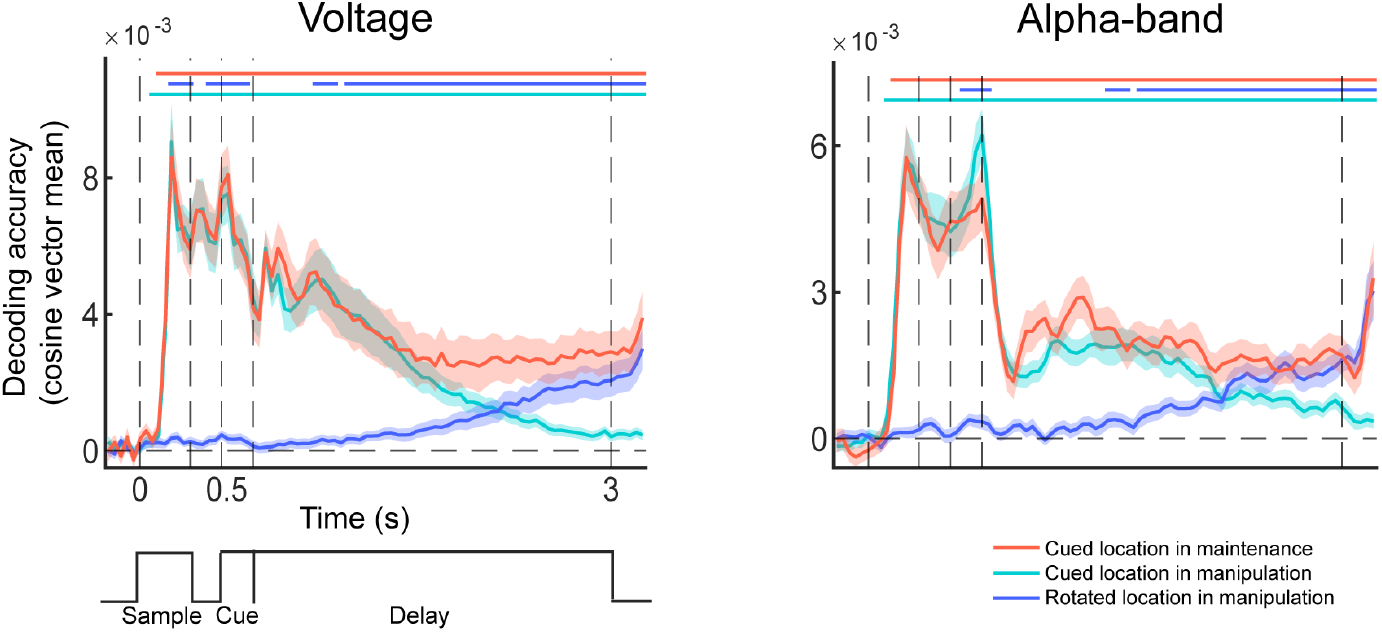
Location decoding from all channels in voltage and alpha-band activity in memory maintenance and manipulation (N=49). Time course of within-condition location decoding results in voltage (left) and alpha-band (right) signals, for cued location in maintenance (red), cued location in manipulation (cyan), and rotated location in manipulation (blue), respectively. Horizontal lines on top denote significance of the corresponding condition.

Having observed robust location representation in memory maintenance and manipulation in both voltage and alpha-band activity, we further investigated potential differences in the spatiotemporal patterns of location representations between these two types of signals. To ensure a fair comparison between neural representations and to boost signal-to-noise ratio, we employed a mixed-model decoding approach, whereby a common decoder was trained on all trials of interest, and tested on each condition separately. Specifically, we trained a common decoder using the cued location in maintenance trials and the rotated location in manipulation trials and focused on the comparison between the two.

We divided all EEG channels into three groups: frontal, central, and posterior channels, and performed the decoding analyses in each group of channels (Fig 3). Our results showed that the cued location could be well decoded throughout a trial (Figs 3A-C); however, decoding pattern for the rotated location differed substantially between voltage and alpha-band activity: specifically, in voltage, the rotated location showed the earliest decoding in central channels (starting around 1.16 s), followed by frontal channels (starting around 1.44 s), and minimal level of decoding only towards the end of delay in posterior channels (starting around 2.92 s; Figs 3A-C). Conversely, in alpha-band, the earliest decoding of the rotated location appeared in posterior channels (starting around 1.48 s), followed by central channels (starting from 1.84 s) and frontal channels (starting around 2.40 s; Figs 3A-C).

**Fig 3.**
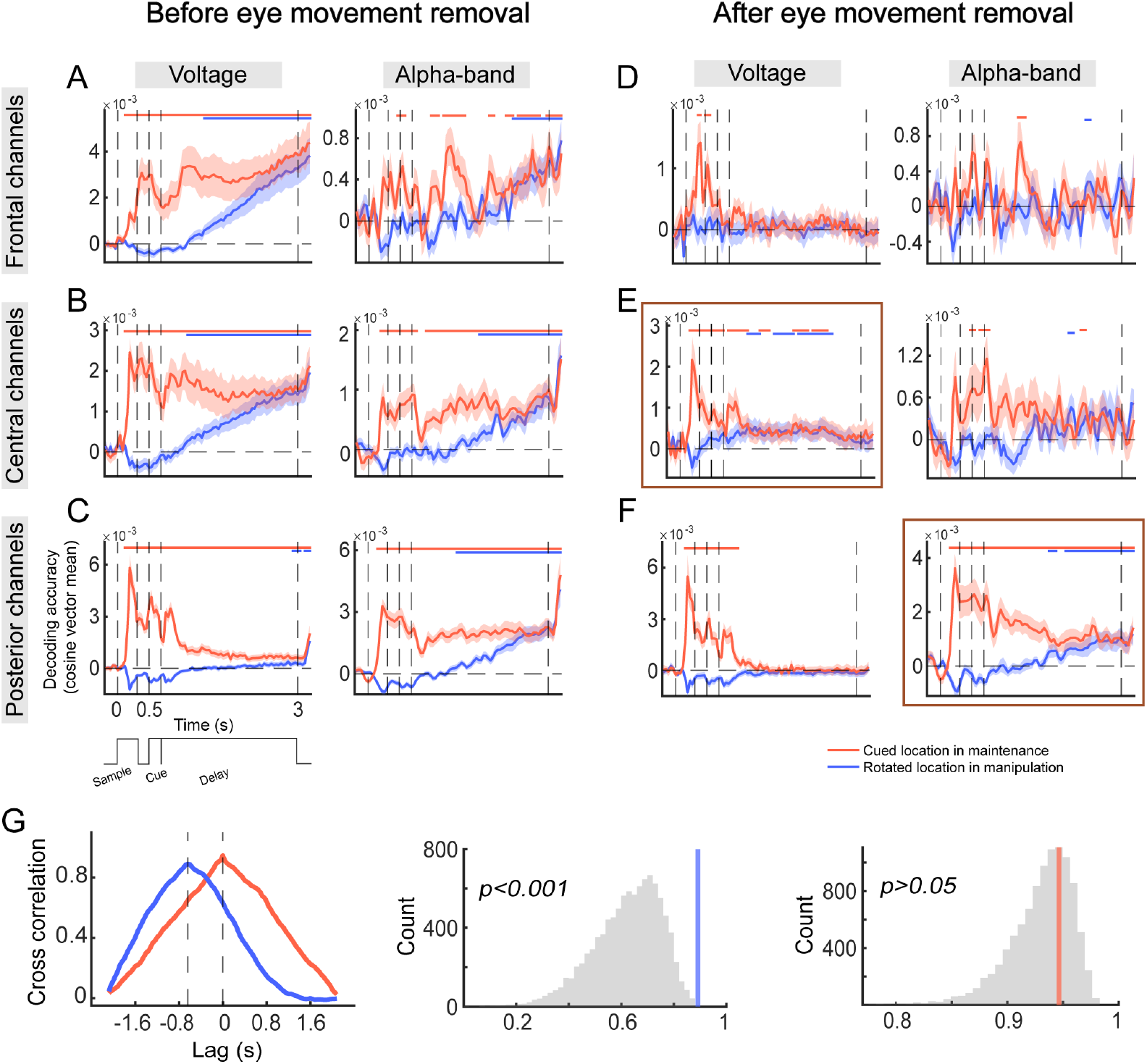
Spatiotemporal dynamics of location representation in voltage and alpha-band activity. (A) Time course of mixed-model decoding results for cued location in maintenance condition (red) and rotated location in manipulation condition (blue) before removing influences of eye movements (N=49). Horizontal lines on top denote significance of the corresponding condition. The left panel displays voltage decoding in frontal channels, and the right panel displays alpha-band decoding in frontal channels. (B) Same conventions as A, but with results from central channels. (C) Same conventions as A, but with results from posterior channels. (D) Time course of mixed-model decoding results for cued location in maintenance condition and rotated location in manipulation condition after removing influences of eye movements (N=23). The left panel displays voltage decoding in frontal channels, and the right panel displays alpha-band decoding in frontal channels. (E) Same conventions as D, but with results from central channels. (F) Same conventions as D, but with results from posterior channels. (G) Cross-correlation between central voltage and posterior alpha-band signals. Blue line represents cross-correlation between decoding time courses for rotated location in manipulation in central voltage and posterior alpha, and red line represents cross-correlation between decoding time courses for cued location in maintenance in central voltage and posterior alpha. Data were obtained from the decoding time courses highlighted by the solid brown boxes in E and F. The left panel shows the cross-correlation coefficient curve, where the x-axis represents the shifted time points of alpha-band time course relative to the central voltage time course (one time point represents 40 ms). The black dashed line indicates the corresponding time lag when each curve reaches the maximum cross-correlation coefficient. The time lag was 640 ms in manipulation and 0 ms in maintenance. The middle and right panels show significance testing of the time lag in each condition. The histogram represents the null distribution of cross-correlation in each condition, and the red and blue vertical lines represent the true maximum correlation coefficient.

However, it is known that spatial-related decoding in EEG can be susceptible to influences from eye movements [17,18], particularly in frontal channels. In the current study, a subset of participants (Experiment 2, N = 23) had been strictly monitored with their eye movements alongside with EEG recording. We thus removed potential influences of eye movements from EEG data by using linear regression to regress out eye gaze effects. We then repeated the mixed-model decoding analyses using the regressed EEG data (Figs 3D-F). Interestingly, results in different groups of channels as well as in voltage and alpha-band activity again displayed differential sensitivity to influences of eye movements: specifically, in frontal channels, all decoding results in delay period disappeared, indicating that decoding in frontal channels was potentially contaminated by eye movements in both types of signals (Fig 3D). For central and posterior channels, results further diverged between voltage and alpha-band activity: for alpha-band activity, sustained decoding of both cued and rotated locations during delay after regressing out eye movements was predominantly observed in posterior channels, suggesting that posterior channels might primarily contribute to alpha-band decoding (Fig 3F). Unlike alpha-band activity, for voltage activity, decoding was only evident during the sample period for the cued location in posterior channels. Interestingly, decoding of both cued and rotated locations remained significant during delay in central channels. This suggests that voltage activity in central but not posterior channels likely contained neural representations of manipulated information independent of eye movements (Fig 3E). This difference in spatial configuration was further confirmed by a decoding searchlight analysis, which revealed that channels contributed to location decoding in voltage activity were mainly located centrally, and those contributed primarily to location decoding in alpha-band activity were located posteriorly (S1C-D Fig). In addition, we demonstrated that decoding results in central voltage were preserved at lower frequency (< 3 Hz) but not at higher frequency bands (3-45 Hz), suggesting distinct signal origins in central voltage as opposed to alpha-band activity (S1A-B Fig).

Besides differences in spatial configuration, we also observed that significant decoding of rotated location in central voltage (around 1.12 s; Fig 3E) emerged earlier in time than that in posterior alpha-band activity (around 1.80 s; Fig 3F). To quantify this potential difference in timing, we calculated the cross-correlation between the two signal time series; indeed, the decoding time course of central voltage preceded that of posterior alpha by approximately 640 ms. As a comparison, no significant time lag was found between the decoding time courses for cued location in maintenance in the two signals (Fig 3G). Together, these results suggested a possible functional discrepancy between central voltage and posterior alpha in memory manipulation: location representation generated internally via memory manipulation might originate from central voltage, while representation of rotated location in posterior alpha might mainly reflect feedback signals.

It is important to note that although we demonstrated strong eye-movement-related effects on location decoding, we lacked causal evidence between the two. In fact, it was also possible that eye movements and EEG signals co-varied, and eye movements might partially reflect the mental processes related to memory maintenance and manipulation. As a final check for this question, we calculated the correlation between the mixed-model decoding results before and after regression. We observed a high correlation in the decoding of rotated location between the two, both in central voltage and posterior alpha-band activity, suggesting that eye movements mainly impacted the magnitude of decoding, rather than the presence of significant decoding, in the two types of signals (S2 Fig).

To summarize, by systematically examining the nature of location representations in voltage and alpha-band activity, here we demonstrated rotated representations in memory manipulation were primarily encoded in voltage signals located centrally and in alpha-band signals located posteriorly. Moreover, rotated representation in central voltage led that in posterior alpha in time, suggesting rotated representation in central voltage might serve as the source of manipulation signals in working memory. Meanwhile, although both types of signals retained rotated representation regardless of whether eye movement influences were removed or not, location representations in alpha-band demonstrated stronger independence of eye movement signals, indicating differential sensitivity of these two types of signals to eye movements.

### Generalization of location representation between memory maintenance and manipulation in both voltage and alpha-band activity

Next, we asked whether location representations in maintenance and manipulation were generalizable to each other, by training the decoder on data from one condition, and tested it on data from the other condition. Because location representations were most evident in central voltage and posterior alpha, hereafter we focused all our primary analyses on these two types of signals. We first performed the generalization analyses without removing eye movement effects, like what previous work did [14,15]. We found that representation of the cued location in the maintenance condition and that of the rotated location in the manipulation condition were highly generalizable to each other, starting around the time point at which the rotated location became decodable; representations of the cued location in the two conditions were also highly generalizable to each other in the early portion of the trial, but became progressively more distinct and even ungeneralizable towards the end of delay. Likewise, representations of the cued location and rotated location in the manipulation condition also became dissimilar as trial proceeded. These patterns were observed in both central voltage and posterior alpha-band activity (Figs 4A-B), but not in posterior voltage wherein rotated representation was absent (Fig 4C). We further confirmed that this divergence in neural codes cannot be explained by passive decay of sensory or mnemonic traces, because in trials with an uncued location (the two-dot trials), representation of the uncued location quickly dropped to baseline after retrocue (Fig 4), at stark contrast with the cued location in manipulation.

**Fig 4.**
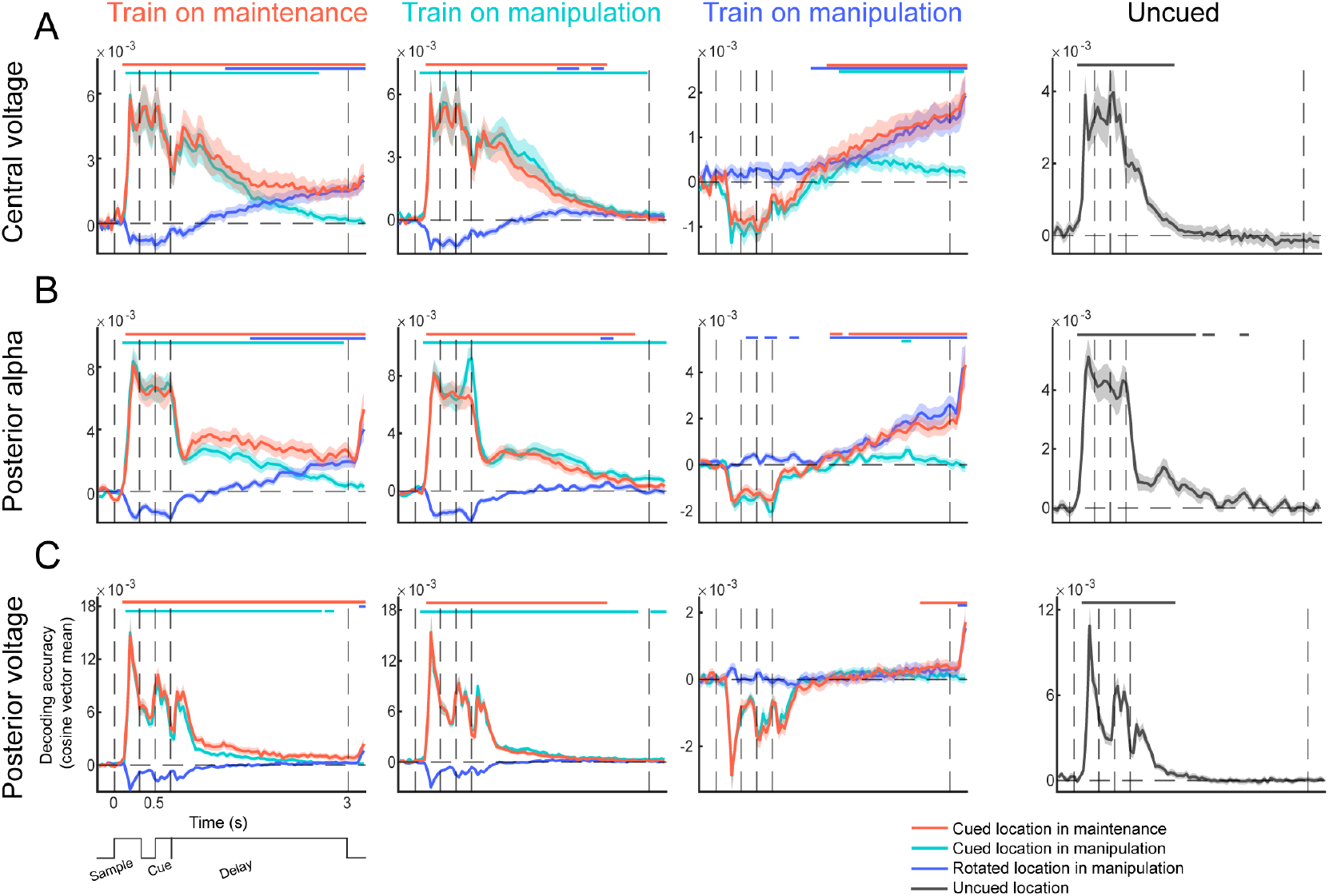
Generalization of decoding results in central voltage, posterior alpha-band and posterior voltage (N=49). (A) Generalization of location decoding between maintenance and manipulation in central voltage activity. From left to right: results from the decoder trained on the cued location in maintenance, and tested on all three conditions; results from the decoder trained on the cued location in manipulation, and tested on all three conditions; results from the decoder trained on the rotated location in manipulation, and tested on all three conditions; result for uncued location. Horizontal lines on top denote significance of the corresponding condition. (B). Same conventions as A, but with generalization results from posterior alpha-band activity. (C) Same conventions as A, but with generalization results from posterior voltage activity.

We also conducted the generalization analyses using the eye-movement-regressed EEG data. Overall, similar generalization patterns remained in posterior alpha-band activity, suggesting generalization of neural codes between maintenance and manipulation in alpha-band activity was less prone to influences of eye movements. In comparison, generalization pattern in central voltage became statistically worse (S3 Fig). Nevertheless, we note that the weaker generalization in central voltage could be partially attributed to a lack of statistical power in the generalization analyses, because when all trials (instead of trials from a single condition) were used to train the decoder as in the mixed-model approach (Fig 3E), both the cued location in maintenance and rotated location in manipulation revealed significant representations in central voltage. These results together suggested shared location representations between memory maintenance and manipulation, and also a gradual separation in neural codes between cued and rotated representations during memory manipulation.

### Location representations as characterized by high-dimensional neural subspaces

Having observed a separation in neural codes between the two location representations in memory manipulation, we next sought to understand how this transformation could underlie memory behavior. Specifically, because participants simultaneously held representations of the cued and rotated locations in the manipulation condition, these two representations could potentially interfere with each other, especially when the rotated representation initially emerged. We therefore hypothesized that neural separation between the two representations was implemented to resist interference; and if so, the degree of separation between the two representations should predict memory performance of the rotated location (Fig 5A). On the other hand, although decoding analysis offers a convenient and powerful tool to understand neural representations of memory maintenance and manipulation, it does not provide a quantified description of the differences between two neural representations. For instance, successful generalization would indicate shared neural representations between cued and rotated locations, but it does not speak to how similar these two representations are. Likewise, failure in generalization would indicate the neural representations of two conditions are dissimilar, but again, it remains unknown how different the representations can be. Here, inspired by recent advances in dimension reduction and neural state space analyses, we investigated the dimensionality of location representations in different conditions as well as in different types of signals, using Principal Component Analysis (PCA) in combination with PCA-based decoding, and quantified the differences between representational subspaces of cued and rotated representations using principal angles [19,20].

**Fig 5.**
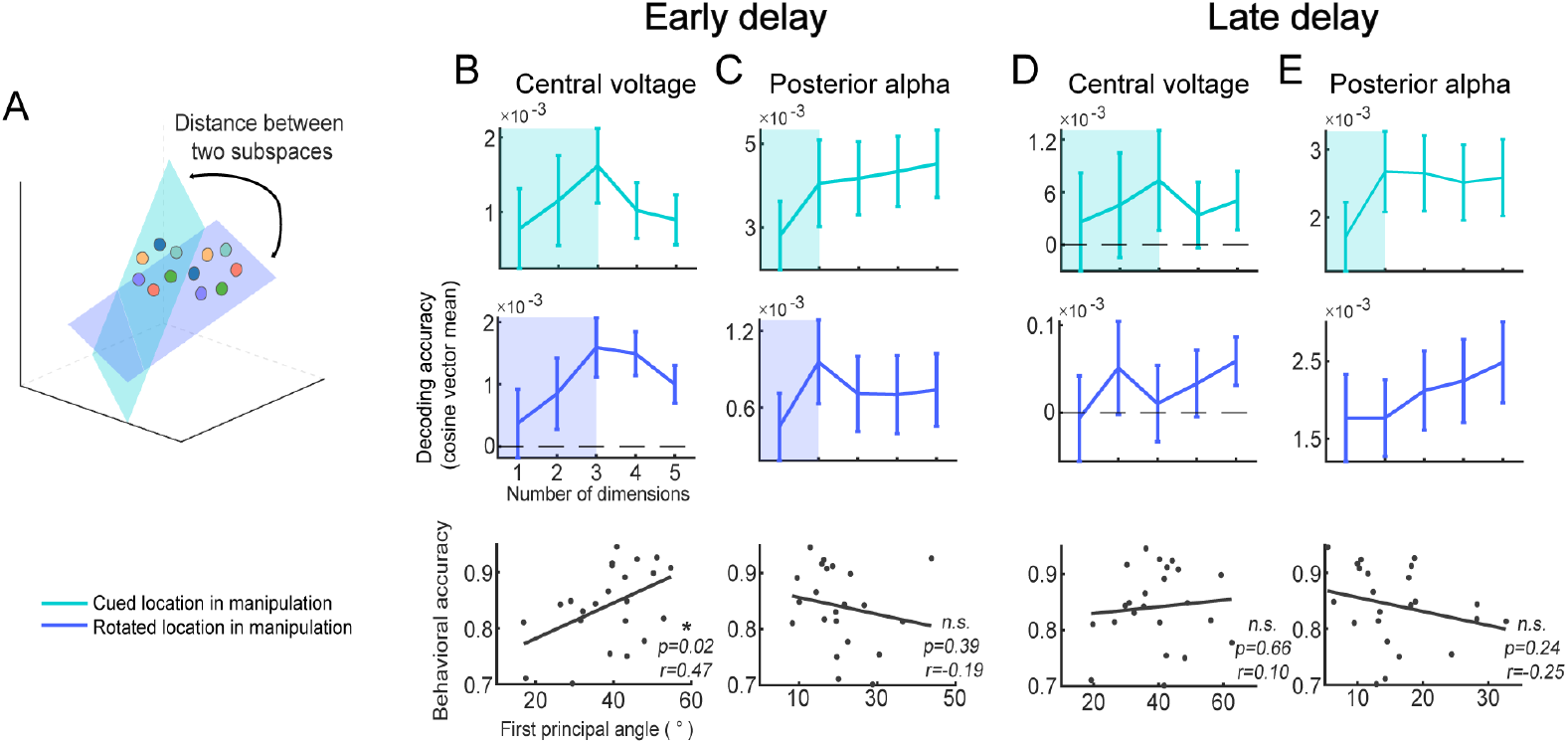
Characterization of neural subspace of the cued and rotated locations in the manipulation condition (after removing influences of eye movements, N = 23). (A) Schematic illustration of the results. The cued and rotated locations are represented in different neural subspaces, indicated by different colors (cyan and blue). The distance between the two subspaces is characterized using the first principal angle between subspaces. (B) Changes in decoding accuracy in central voltage with increasing subspace dimension in the early delay, for the cued location (top panel) and rotated location (middle panel) in manipulation, respectively. Shaded area indicates the dimension of the subspace for the corresponding location. Both the cued and rotated locations are three-dimensional. Bottom panel illustrates the correlation between the first principal angle of the two subspaces and individual participant’s behavioral accuracy. (C) Same conventions as B, but with results from posterior alpha-band in the early delay. (D) Same conventions as B, but with results from central voltage in the late delay. E. Same conventions as B, but with results from posterior alpha-band in the late delay.

We defined the dimensionality of location representation as the minimal number of PCs required to reach ceiling performance in location decoding. To facilitate comparison, we performed the dimensionality analysis in two pre-defined time periods: early and late delay. We observed that in early delay during which the rotated representation started to emerge, both the cued and rotated locations in the manipulation condition were represented in a three-dimensional subspace in central voltage (Fig 5B), and were represented in a two-dimensional subspace in posterior alpha (Fig 5C). This indicated that although both central voltage and posterior alpha-band activity exhibited location representations, the dimensionality of location representation in central voltage was higher. This result was further confirmed by the cumulative explained variance of the PCs, which showed that posterior alpha-band activity required fewer PCs to reach a cumulative explained variance of around 85% (S4 Fig).

Having identified the neural representational subspaces of cued and rotated locations, we next asked whether the subspaces could provide a quantified description of the differences between locations. We found that the distance between the representational subspaces of the rotated and the cued location in central voltage was positively correlated with participants’ behavioral accuracy (*r* = 0.47, *p* = 0.02; Fig 5B). In other words, larger separation between subspaces was associated with better behavioral performance on memory manipulation. By contrast, we did not observe a correlation with behavior in posterior alpha-band activity (*r* = -0.19, *p* = 0.39; Fig 5C).

When applying the same analysis to the late delay, we found that cued location was also represented in a three-dimensional subspace in central voltage and in a two-dimensional subspace in posterior alpha. However, the rotated location showed higher dimensionality in both central voltage and in posterior alpha, probably due to decreased decoding performance and increased noise (Figs 5D-E). Although we were unable to determine their dimensions precisely, for completeness, we calculated the principal angle using the same dimensionality as the cued locations, but did not observe significant correlation with behavior in either type of signals (central voltage: *r* = 0.10, *p* = 0.66; posterior alpha: *r* = -0.25, *p* = 0.24), suggesting that early delay was the critical time period in determining memory performance.

Lastly, we examined whether decoding accuracy alone was related to behavioral performance (S5 Fig). In central voltage, there was a significant correlation between decoding accuracy and participants’ reaction time during the sample and early delay periods for the cued location in the maintenance condition (S5A Fig). However, no significant correlation was observed for late delay, for the manipulation condition, or for any decoding in posterior alpha-band activity (*p*s > 0.10). In other words, during memory maintenance where no separation in neural codes was required, the strength of location representation in central voltage was also predictive of memory behavior.

## Discussion

In the present study, we set out to investigate the neural signatures of stimulus representation in memory maintenance and manipulation. Leveraging EEG and multivariate decoding, we demonstrated robust neural representations of maintained and manipulated information in both voltage and alpha-band activity. However, these two types of signals displayed dissociable spatiotemporal dynamics in memory manipulation: manipulated representation was most evident in voltage activity in central channels, and in alpha-band activity in posterior channels. Moreover, the temporal emergence of manipulated representation in central voltage preceded that in posterior alpha, suggesting central voltage might serve as the source of manipulated information in working memory. Location representation was also higher-dimensional in central voltage compared to posterior alpha, and location representation in central voltage exclusively predicted multifaceted memory behavior. Altogether, these results provide a comprehensive understanding of the functional distinction between voltage and alpha-band activity in memory manipulation, from both the spatiotemporal and representational perspectives.

Extant work on working memory has revealed significant stimulus representations in both voltage and alpha-band activity during memory delay [2–7]. Nevertheless, the exact functions of these two types of signals in working memory have remained controversial. Specifically, because maintained information was always the same as sensory sample information during memory maintenance, it remains unclear whether stimulus representations, especially those observed in posterior channels in EEG, were merely a reflection of passive sensory traces persisted or reactivated during memory delay, rather than mnemonic signals generated internally to sustain memory. The answer to this question would also be crucial for theories of working memory, because the neural locus of memory generation and maintenance has been central to the debates in the field of working memory over the past decade [21,22]. Here, we tackled on this problem by systematically examining the contribution of voltage and alpha-band activity to memory manipulation. Unlike memory maintenance, memory manipulation requires the generation of mnemonic information from an internal source without being contaminated by sensory signals. We observed several distinct characteristics of voltage and alpha-band activity in memory manipulation: first, their spatial layout was different: location representation in voltage activity was most prominent in channels located centrally, and location representation in alpha-band activity was most prominent in channels located posteriorly; Second, their temporal dynamics were different: location representation in central voltage significantly preceded that in posterior alpha in time; third, their representational dimensionality was different: location representation in central voltage was higher-dimensional than that in posterior alpha; lastly, their behavioral predictability was different: location representation in central voltage exclusively predicted memory behavior, in terms of both location decoding accuracy and subspace separation. Collectively, these results suggest distinct functions of voltage and alpha-band activity in memory manipulation, and highlight the role of central voltage in the generation of information in memory manipulation.

Further analyses demonstrated that location representation in central voltage primarily originated from lower frequency signals below 3 Hz. This low-frequency nature of the signals would be consistent with recent observation in the perceptual awareness literature, which showed that slow cortical potentials below 5 Hz encoded perceptual contents during bistable perception, while alpha-band activity did not [23]. On the other hand, the critical role of central voltage in memory manipulation does not deny the importance of posterior alpha-band activity in working memory. In fact, we found that location representation in posterior alpha-band activity was more persistent than that in central voltage, especially in memory maintenance. It also displayed higher robustness to influences from eye movements. Therefore, we interpret stimulus representation in posterior alpha as reflecting feedback signals from higher-order cortex that contribute to the consolidation and rehearsal of stimulus information in working memory, consistent with the sensory recruitment hypothesis of working memory [21], as well as the layer-specific fMRI finding in early visual cortex [13]. It is noteworthy that the current observation of the spatiotemporal distinction between central voltage and posterior alpha is also broadly in line with another recent work from our lab on the spatiotemporal dynamics of self-generated imagery [24].

Besides these spatiotemporal characteristics of central voltage and posterior alpha-band activity, we further demonstrated a clear functional dissociation between the to-be-responded (i.e., cued location in maintenance and rotated location in manipulation) and the not-to-be-responded (i.e., cued location in manipulation) locations. Specifically, we found that neural representations of cued location in maintenance and rotated location in manipulation, regardless of the specific task condition, were highly similar; while neural representation of cued location in manipulation diverged from the other two towards the end of delay. This observation is consistent with the recent theoretical proposal of action-oriented working memory, which posits that memory contents are selected and represented to guide future actions and behavior [25–27]. Notably, the action-unrelated, cued location in manipulation was still represented actively, although in a different format from the action-related locations. This active representation was distinct from representation of the uncued location, which quickly dropped to baseline after retrocue, suggesting another dissociation between the action-unrelated yet task-relevant and the task-irrelevant neural representations in working memory. In other words, the action-unrelated yet task-relevant information underwent a task-driven recoding from the action-related information probably to resist interference, similar to the priority-based recoding as observed in attentional prioritization studies [28,29]. In light of these findings, we characterized the separation between the action-related and action-unrelated neural codes by applying a novel PCA-based decoding and dimension reduction approach to EEG data to calculate the dimensionality of stimulus representation and constructed low-dimensional stimulus subspace in both types of signals. This approach provided a quantified description of the neural similarity between location representations beyond classic cross-condition generalization methods. Specifically, the degree of neural similarity or separation between representations can be quantified by the distance between neural subspaces of stimuli. Results from this approach further revealed that although stimulus representation in both types of signals could be compressed into a low-dimensional neural state space, only the degree of separation between cued and rotated locations in central voltage was predictive of subsequent memory performance on rotation. This result provided further support that stimulus representation in central voltage is critical for memory manipulation.

Our results that stimulus information was primarily represented in central but not posterior voltage activity might seem at odds with previous work at first glance. However, many previous studies with significant stimulus decoding in posterior voltage activity have used a visual impulse approach. In those studies, decoding of stimulus information quickly decayed after stimulus offset; by presenting a visual impulse in the middle of the delay, the stimulus information in posterior voltage could be reactivated for a brief period of time [5,9,30]. These results have been taken as evidence for “activity-silent” neural codes in working memory that was otherwise undetectable in the absence of visual impulses. In our case, successful decoding of rotated location was observed in central voltage without the presence of visual impulses, whereas rotated representation in posterior voltage was largely absent as in previous work [15]. This suggested that stimulus representations in posterior and central voltage might have inherently distinct natures, with stimulus representation in posterior voltage being in an “activity-silent” mode, and that in central voltage being in an active, persistent mode. Previous fMRI work demonstrated that the lateral prefrontal cortex (lPFC) showed higher persistent neural activity in memory manipulation than in memory maintenance, suggesting that lPFC might be critical for memory manipulation [31,32]. It was thus possible that central voltage reflects these manipulation-related signals arising from frontal cortex. Future studies using techniques with higher spatial resolution such as magnetoencephalography (MEG) may provide further insights into the separate sources of central and posterior voltage signals.

To conclude, we identified two important neural signatures, central voltage and posterior alpha-band activity, in representing stimulus information in memory manipulation. We propose that both neural signatures are important for working memory; yet, central voltage activity might serve as the origin of internally-generated, manipulated information, and is closely linked to memory behavior. By contrast, posterior alpha-band activity might be crucial for the sustained maintenance of stimulus information, potentially received from top-down feedback signals. Future work on working memory that simultaneously focuses on these two neural signatures will help to gain a more coherent understanding of the neural mechanism underlying working memory.

## Methods

### Participants

Fifty adults were recruited from the community of Shanghai Institutes for Biological Sciences, Chinese Academy of Sciences, with 26 participants in Experiment 1 and 24 participants in Experiment 2. One participant was excluded from Experiment 2 due to malfunction of the EEG equipment. Therefore, the final analysis included 26 participants in Experiment 1 (11 males, mean age = 23.1±1.1 years), and 23 participants in Experiment 2 (6 males, mean age = 24.1±2.0 years). All participants had normal or corrected-to-normal vision and normal color vision, and reported neurologically healthy. All participants gave written informed consent approved by the ethical committee of Center for Excellence in Brain Science and Intelligence Technology, Chinese Academy of Sciences (CEBSIT-2020028), and were monetarily compensated for their time.

### Apparatus and stimuli

Experimental stimuli were generated and controlled by MATLAB (R2018b, The Mathworks) and Psychtoolbox-3 extensions [33]. Visual stimuli were presented on a 23-inch (58.42 cm) screen with a refresh rate of 60 Hz and a resolution of 1280 × 720. Viewing distance was set at 60 cm. Participants completed the task with a computer mouse. The background of the screen remained grey (RGB = 128, 128, 128) throughout the experiment, and a fixation cross (∼0.6° of visual angle) was presented at the center of the screen throughout the experiment. Fixation cross was white during task period and turned black during Inter-trial Intervals (ITIs). Stimuli were either red or blue dots (∼0.6° of visual angle) presented at a visual angle of 3.7° from the center of the screen, and were selected from 6 bins (16°, 76°, 136°, 196°, 256° and 316°) with a random jitter of ± 10° in each bin.

### Procedure

Participants completed a spatial working memory task consisting of two conditions: maintenance and manipulation. On each trial, participants viewed either one or two sample dots presented at different spatial locations at the beginning of the trial. A retrocue followed to indicate the color of the to-be-maintained or manipulated item. In the maintenance condition, participants were required to maintain the cued dot throughout the trial; In the manipulation condition, participants were required to mentally rotate the cued dot following a rotation cue.

Each trial began with presentation of the sample dot(s) for 300 ms. For one-dot trials, a single sample dot was randomly selected from the six bins, and the color of the dot was either blue or red. For two-dot trials, the two sample dots were sampled independently from the six bins without replacement, and the colors of the two dots were blue and red, respectively. After an interval of 200 ms, a digit cue (1° of visual angle) was shown for 200 ms at the center of the screen. The color of the cue indicated the color of the to-be-maintained or manipulated sample dot; specifically, for one-dot trials, the color of the cue was always the same as the color of the sample dot. The numerical value of the cue indicated the angle by which the participants needed to rotate the sample dot. Specifically, in the maintenance condition, the digit was always 0. In the manipulation condition, the digit was randomly selected from ±60, ±120, or 180 (with positive values indicating clockwise rotation and negative values indicating counterclockwise rotation). A delay period of 2300 ms followed the disappearance of the cue. During the delay period, the participants needed to maintain the original location of the cued dot in the maintenance condition, or to mentally rotate the cued dot by the specific angle in the manipulation condition. At the end of delay, a response screen appeared, consisting of six probes arranged in a 2×3 matrix at the center of the screen as shown in Fig 1A. Each probe consisted of a circle (2.4° of visual angle) with a dot (0.35° of visual angle) on it. One of the probes contained the correct location of the maintained or rotated dot, and dot locations in the five other probes were ±60°, ±120°, or 180° from the correct location. Participants were unaware of the configuration of the probes, and were only told to select the probe with the correct dot location. Response period ended when participants entered their response using the left mouse button, or after a maximum response time of 2300 ms. ITI varied randomly between 600 and 900 ms.

Experiment 1 and Experiment 2 shared the same task procedure, differing only in the number of trials participants were required to complete. In Experiment 1, each run consisted of 26 maintenance trials (12 one-dot trials and 14 two-dot trials) and 52 manipulation trials (24 one-dot trials and 28 two-dot trials). In total, participants completed 18 runs, resulting in 468 maintenance trials and 936 manipulation trials. In Experiment 2, each run consisted of 28 maintenance trials (14 one-dot trials and 14 two-dot trials) and 42 manipulation trials (20 one-dot trials and 22 two-dot trials). In total, participants completed 18 runs, resulting in 504 maintenance trials and 756 manipulation trials. In both experiments, all trials were interleaved in a randomized order. We chose to collect more trials for the manipulation condition given that multiple rotation angles were used in this condition. Moreover, because the primary goal of the current study was to investigate the neural mechanism underlying working memory maintenance and manipulation, we combined one-dot and two-dot trials within each condition for all subsequent analyses (except for decoding of uncued locations). Trial number within each condition was equated for subsequent analyses when needed. Prior to the main task, each participant underwent a practice session on the task, which ended when the participant reached an average behavioral accuracy of above 80%.

### EEG acquisition and preprocessing

EEG was recorded from 64 electrodes (Brain Products actiCHamp), laid out according to the international 10–10 system. Data were recorded at 1000 Hz using a Brain Vision amplifier and Recorder (Brain Products GmbH, Gilching, Germany). The midline central electrode (FCz) was attached to the forehead as the reference. In Experiment 2, four electrodes (FT9, FT10, TP9 and TP10) were used for electrooculography (EOG), two were placed ∼1-2 cm above and below the left eye or right eye, and two were placed ∼1-2 cm lateral to the external canthi. In Experiment 1, the same four electrodes were discarded. The impedances of all electrodes were kept below 30 kΩ.

EEG preprocessing was conducted using EEGLAB [34] and custom codes. Raw data were downsampled to 250 Hz and bandpass filtered (0.01-45 Hz). For all electrodes (a total of 59 electrodes, excluding four electrodes for EOG and one for reference), the data were epoched relative to trial onset (-500 ms to 3500 ms). Epochs with voltage drifts, muscle and eye movement artifacts were identified visually and removed from all subsequent analyses. Eye blink artifacts were identified and removed using independent components analysis (ICA). For subsequent analyses, each epoch was baselined using signals from −200 ms to 0 ms before trial onset. We performed time-frequency decomposition by using Morlet wavelet convolution [35]. Voltage data from each channel and trial were convolved with a family of complex Morlet wavelets spanning 3–45 Hz in 1 Hz steps with wavelet cycles increasing linearly between 3 and 10 cycles as a function of frequency. Power in each frequency was calculated as the percent change of squared absolute value in the resulting complex time series relative to the baseline between −200 ms and 0 ms.

To save computation time, voltage and power signals were further downsampled to 25 Hz in the decoding analyses. In some analyses, we divided the 59 electrodes into three groups based on their locations. Specifically, Frontal channels included Fp1, Fp2, AF7, AF3, AFz, AF4, AF8, F7, F5, F3, F1, Fz, F2, F4, F6 and F8. Central channels included FT7, FC5, FC3, FC1, FC2, FC4, FC6, FT8, T7, C5, C3, C1, Cz, C2, C4, C6, T8, TP7, CP5, CP3, CP1, CPz, CP2, CP4, CP6 and TP8. Posterior channels included P7, P5, P3, P1, Pz, P2, P4, P6, P8, PO7, PO3, POz, PO4, PO8, O1, Oz and O2.

### Eye-tracking acquisition and preprocessing

In Experiment 2, we additionally monitored eye gaze position using a desk-mounted infrared eye-tracking system (EyeLink 1000 Plus, SR Research, Ontario, Canada) at a sampling rate of 1000 Hz. Two participants used remote mode and the others used chin-rest mode. Eye-tracking data were preprocessed using methods provided in previous work [36] and custom codes. Data were first filtered using a Savitzky-Golay (SG) FIR smoothing filter. Eye blinks and other artifacts were identified and removed using a velocity-based algorithm and acceleration criteria. Missing values caused by eye blinks were filled in using spline interpolation. Data were then baseline corrected using the average data from -200 ms to 0 ms prior to sample onset, and downsampled to 25 Hz.

### Distance-based decoding

To decode different spatial locations in EEG signals, we used a distance-based decoding method [5,9], based on the Mahalanobis distance between the neural responses to different locations. For each condition, we partitioned the data for 8-fold cross-validation using the *cvpartition* function in MATLAB. In each iteration, all trials in the 7 training folds were used to calculate the covariance matrix using a shrinkage estimator [37]. The trial number for each location bin in the training folds was resampled to match that of the bin with the fewest trials. The resampled training trials for each bin were then averaged, and convolved with a half cosine basis function raised to the 5th power. Next, we used *pdist2* function in MATLAB to calculate the Mahalanobis distance between each trial in the testing fold and the averaged and convolved training data for each bin. This process yielded 6 distances for each test trial. Finally, we used the cosine-weighted mean of the distances as the decoding performance of each trial. This procedure was repeated for all iterations, and the results were averaged after 100 repetitions.

We utilized this method for three analyses:

1. Within-condition decoding: We used the data from each condition as both the training and testing set to obtain the decoding accuracy for each condition.
2. Generalization decoding: To investigate whether neural representations between conditions could be generalized, we used data from one condition as the training set and data from the other conditions as the testing set, to test whether neural representation in the training condition could be generalized to the testing condition.
3. Mixed-model decoding. To compare the decoding performance between two conditions, the training set consisted of data from both conditions (mixed-model), and data from each condition were tested separately. This approach ensured that the comparison was based on the same decoder.

Given that the trial number in the maintenance condition was fewer than that in the manipulation condition, we balanced the number of trials for each location bin in each condition in all three decoding methods. To make full use of all data and to obtain reliable estimates, we repeated the subsampling and decoding process 100 times.

We assessed the significance of decoding performance using one-sample t-tests in combination with a cluster-based permutation procedure to correct for multiple comparisons. Specifically, one-sample t-tests were first performed on the decoding results to identify clusters with significant decoding performance. Next, for each participant, the decoding accuracy at each time point was randomly multiplied by either 1 or -1, and one-tailed t-tests were performed on the sign-flipped data. Likewise, clusters with significant decoding were identified. This procedure was repeated 10000 times, and the sum of t-values in the largest cluster in each permutation was taken to create a null distribution of permutated sum of t-values. Finally, the sum of t-values in each cluster in the real data was compared against the null distribution to identify clusters with significance (α=0.01).

### Distance-based decoding (with linear regression)

In Experiment 2, eye gaze data were simultaneously recorded with EEG. We used linear regression to remove potential impacts from eye movements on EEG decoding [18]. Specifically, for each electrode and time point, we fitted the following linear regression model:

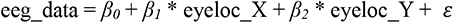

Where eeg_data are the EEG signals, eyeloc_X and eyeloc_Y are the x and y coordinates of participants’ gaze position at the corresponding time point. We used all trials to fit the linear regression. Subsequently, we used the residual of each trial as the input of EEG signal and conducted the distance-based decoding analysis as described above.

### Searchlight decoding analysis

In order to verify the spatial topographical pattern of the decoding results, we performed a searchlight analysis on voltage and alpha-band signals, respectively, on a pre-defined time window. For voltage signals, we used a time window of 1.28 s to 1.84 s. For alpha-band signals, we used a time window of 2.04 s to 2.60 s. The choice of the time windows was based on the fact that voltage signals showed stable decoding early during the delay, while alpha-band signals exhibited relatively stable decoding at a later time. The center of the searchlight moved around all EEG channels. For each electrode and its adjacent electrodes (15 time points × 2-4 neighboring electrodes, the nearest and second nearest electrodes), we used a mixed-model decoding approach, and repeated this process 100 times to make full use of the available data [38,39].

### Cross-correlation

To quantify the timing differences in decoding between different signals, we performed cross-correlation analysis [40] using the *xcorr* function in MATLAB. Specifically, we calculated the cross-correlation between the average decoding performance of the central voltage and posterior alpha-band data from the first decodable time point in the delay period till the end of the epoch. When shifting the latter relative to the former, we calculated the normalized correlation coefficient of these two decoding time series. If there was no timing difference between the two, the maximum correlation coefficient would occur at a lag of 0 ms. If the decoding time course of the central voltage preceded that of posterior alpha, the maximum correlation coefficient would occur at a negative lag. To assess the statistical significance, we compared the maximum correlation coefficient with a null distribution of cross-correlation coefficients. The null distribution was generated by shuffling participant-level signal types (central voltage or posterior alpha), recalculated the mean decoding time course, and repeated the procedure 10000 times.

### Principal component analysis (PCA) and calculation of location-based dimensionality

To better understand location representations in the high-dimensional neural state space, we performed PCA on data for each condition. Specifically, for a given condition, we organized data into a matrix X of size *N_location bins_ *N_electrodes_*, by taking the mean of all trials within each location bin over a specific time period [19]. We focused on two time periods for this analysis, early delay and late delay, by dividing the delay period into two equal segments. Here, *N_location bins_* represents the number of location bins (6 in total), and *N_electrodes_* represents the number of electrodes. After subtracting the column-wise mean of X, we used the *eig* function in MATLAB to perform eigen decomposition on the covariance matrix (X^T^X) and obtained the eigenvector matrix W and the eigenvalue matrix L. The size of W is *N_electrodes_ * N_electrodes_*, with each column representing the weights of each principal component (PC). Therefore, W constructs the transformed space after the corresponding data has been transformed by PCA. L is a diagonal matrix, where the diagonal represents the eigenvalues corresponding to each PC.

Next, we tested the dimensionality of location representation in the neural space constructed by PCA. Dimensionality was defined as the minimal number of PCs required to reach a plateau of location decoding. To achieve this, we first performed PCA-based decoding on data projected onto the PCs obtained from PCA, for each participant separately. Specifically, for each PCA, we obtained a matrix R of size *N_electrodes_ *N_trials_* by concatenating all trials of the corresponding condition for the specific time period (early or late delay). We then multiplied R by the eigenvectors of specific PCs to obtain the projected data matrix P of size *N_trials_*N_pcs_*. For example, to project the neural data onto the three-dimensional subspace constructed by the first three PCs, we multiply R by the first three columns of W (size: *N_electrodes_ *3*), resulting in a P of size *N_trials_*3*. We then performed the distance-based decoding analysis on P, to obtain location decoding performance of the first 3 PCs. To avoid double-dipping in this procedure, for each decoding result, we obtained a null distribution of decoding performance by randomly shuffling the location labels and repeating the above PCA and decoding procedures 1000 times. We took the mean of the null distribution as baseline, and subtracted this baseline from the actual decoding result to obtain the final decoding result.

This PCA-based decoding procedure was repeated iteratively by increasing the number of PCs from 1 to 5 (the maximum number of PCs in this PCA), which resulted in a curve of decoding as the principal components (dimension) increased. We determined the dimensionality based on the increasing rate of the decoding curve: if the decoding accuracy of k+1 PCs exceeded that of k PCs by over 10%, we considered the k+1 th PC to make a significant contribution to decoding. Otherwise, dimensionality of location decoding was set to k.

We also examined the dimensionality of the original data by calculating the cumulative explained variance as the number of PCs increased, without considering their contributions to decoding. For each PCA, using the obtained eigenvalues, we calculated the percentage of explained variance for each PC as *(Val_ipc_ / Val_allpc_) *100*. We then computed the cumulative explained variance as the number of PCs increased.

### Calculation of subspace distance

After determining the dimensionality (k) of location representations in each condition using PCA, we could therefore use the first k PCs to construct the neural subspace of location representations. Specifically, in the manipulation condition, participants simultaneously held representations of the cued and rotated locations during delay, and these two representations could potentially interfere with each other. We therefore hypothesized that the degree of separation between the two representations was related to memory performance of the rotated location. To directly test this idea, we used the principal angle between subspaces [19,20] as a measure of the distance between subspaces. Specifically, we performed singular value decomposition (SVD) on the inner product matrix W_cued_^T^W_rotated_ using the *svd* function in MATLAB. Here, W_cued_ represents the eigenvector matrix corresponding to the subspace of the cued location, and W_rotated_ represents the eigenvector matrix corresponding to the subspace of the rotated location. Taking Fig 5B as an example, the size of both W_cued_ and W_rotated_ was *N_electrodes_*3*. If the dimensions of the two subspaces were inconsistent, we chose the lower dimension of the two as the dimension of the matrix used to calculate the principal angle. For instance, in Fig 5D, we took the first two dimensions of the two subspaces to calculate the principal angle. The SVD resulted in a diagonal matrix where the values on the diagonal were the cosine values of the principal angles. We used the first principal angle along the diagonal to calculate the Pearson correlation between the principal angle and behavioral performance across participants.

### Correlation analysis

Pearson correlations at the participant level were evaluated using the *corr* function in MATLAB. Specifically, to examine the relationship between behavioral performance and neural results, we conducted correlation analyses between behavioral performance (including accuracy and RT) and decoding accuracy, as well as between behavioral performance and the first principal angle. To examine the consistency of decoding results before and after removing the effect of eye movement, we also performed correlation analyses on the decoding time course before and after applying linear regression, and we used a cluster-based permutation method to correct for multiple comparisons across time. Specifically, individual correlation values were shuffled within condition (before or after linear regression), and Pearson correlation was calculated for the shuffled data. This procedure was repeated 10000 times. Clusters with the largest cluster size in each iteration were selected to create the null distribution.

## Acknowledgements

This work was supported by the Ministry of Science and Technology of China (STI2030-Major Projects 2021ZD0203701, 2021ZD0204202), the National Natural Science Foundation of China (32271089), Shanghai Pujiang Program (22PJ1414400), CAS Project for Young Scientists in Basic Research (YSBR-071), and Shanghai Municipal Science and Technology Major Project (2018SHZDZX05).

---

**S1 Fig.**
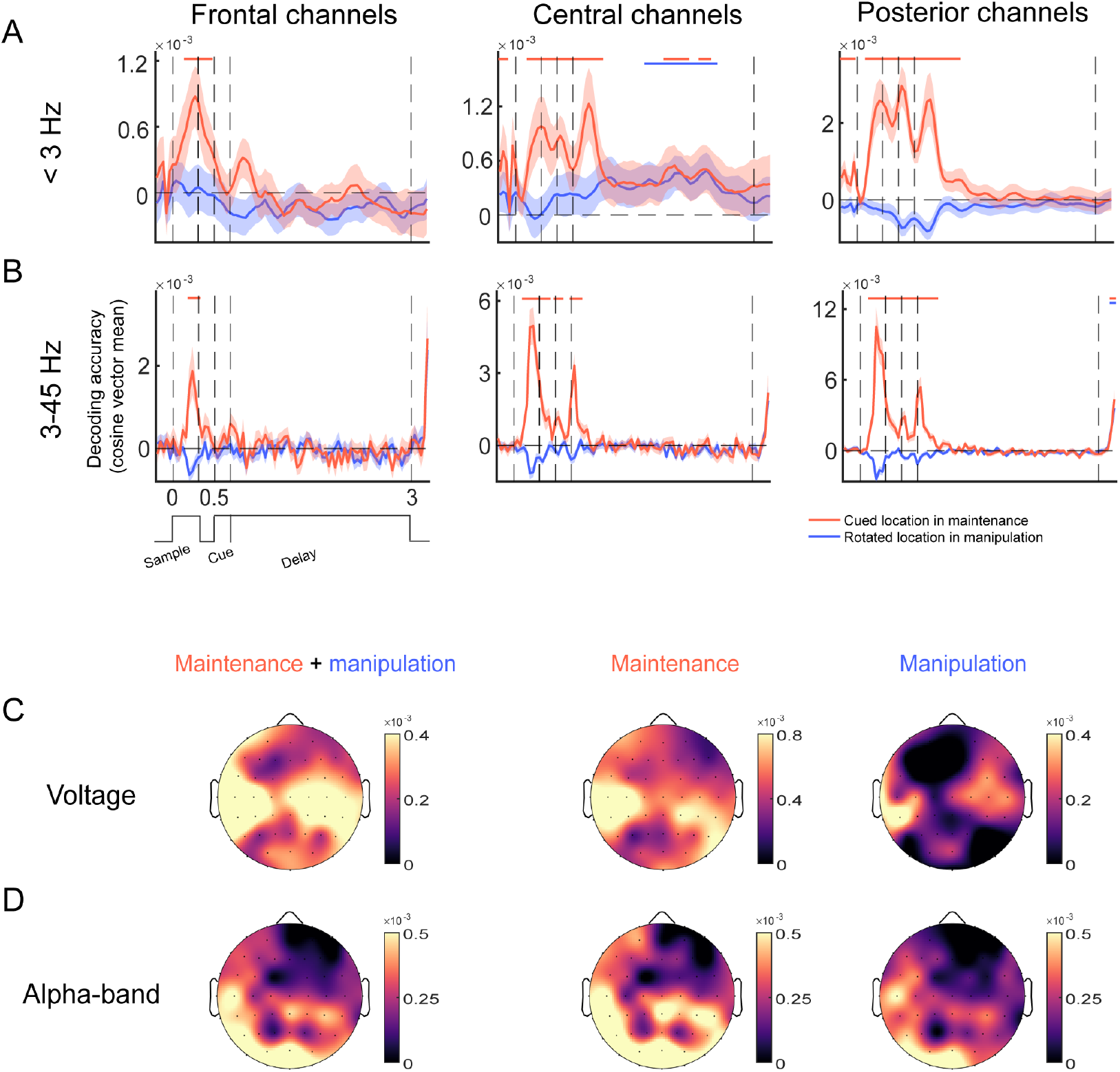
Frequency-specific and spatial localization of decoding results (after removing influences of eye movements, N=23). (A-B) Mixed-model decoding results for voltage signals below 3 Hz (A) and between 3-45 Hz (B) in frontal, central, and posterior channels, respectively. (C-D) Topographic map for mixed-model searchlight decoding results for voltage (C) and alpha-band (D) activity. The left panel represents the average searchlight result of cued location in maintenance and rotated location in manipulation. The middle and right panels represent the searchlight result of the cued location in maintenance and the rotated location in manipulation, respectively. Colorbar denotes decoding accuracy.

**S2 Fig.**
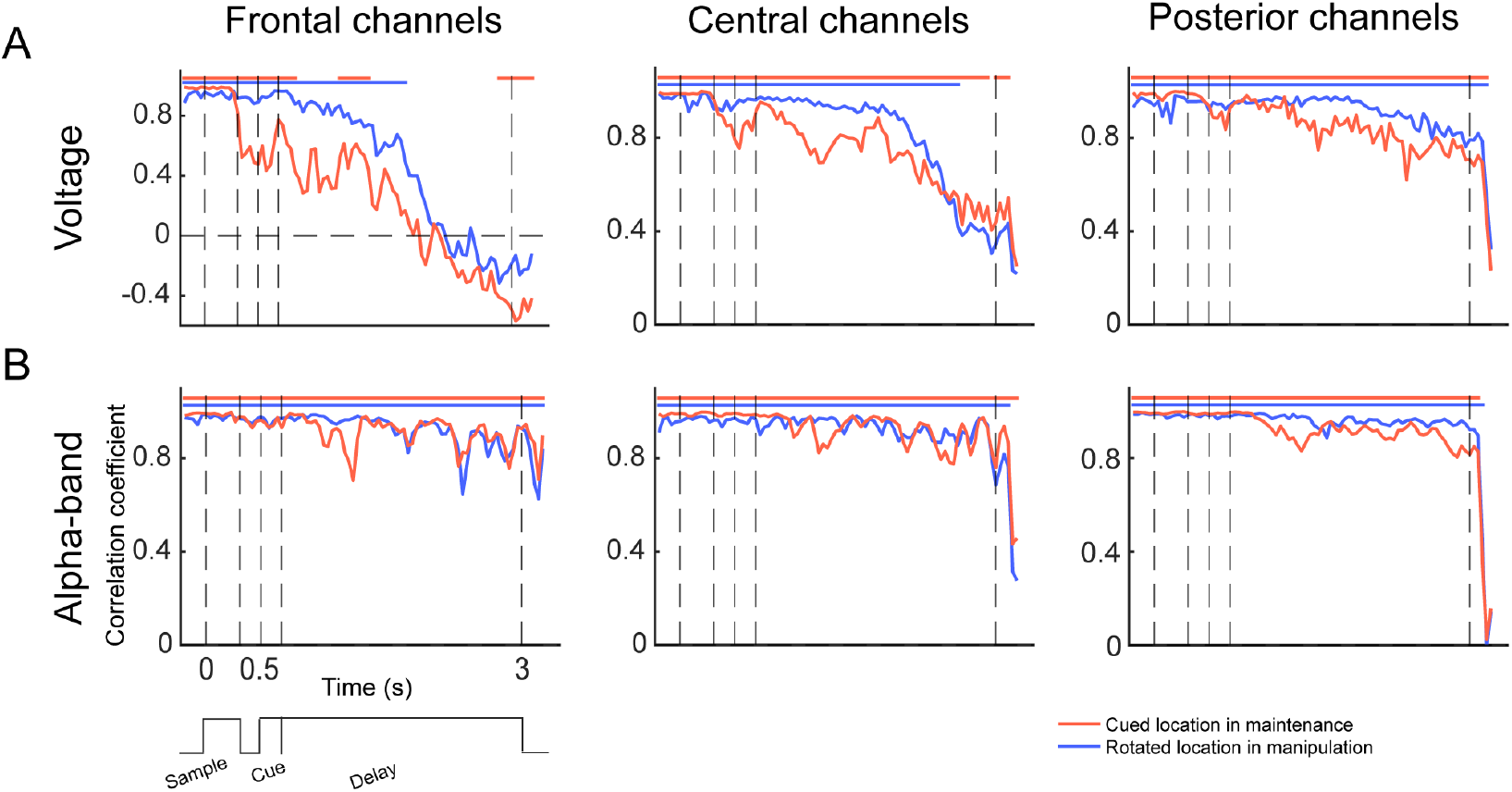
Pearson correlations between decoding results before and after removing eye movement effects (N = 23). (A) Correlation between the mixed-model decoding time courses before and after removing the influence of eye movements, for voltage results in frontal, central, and posterior channels, respectively. (B) Same conventions as A, but with alpha-band results in frontal, central, and posterior channels, respectively.

**S3 Fig.**
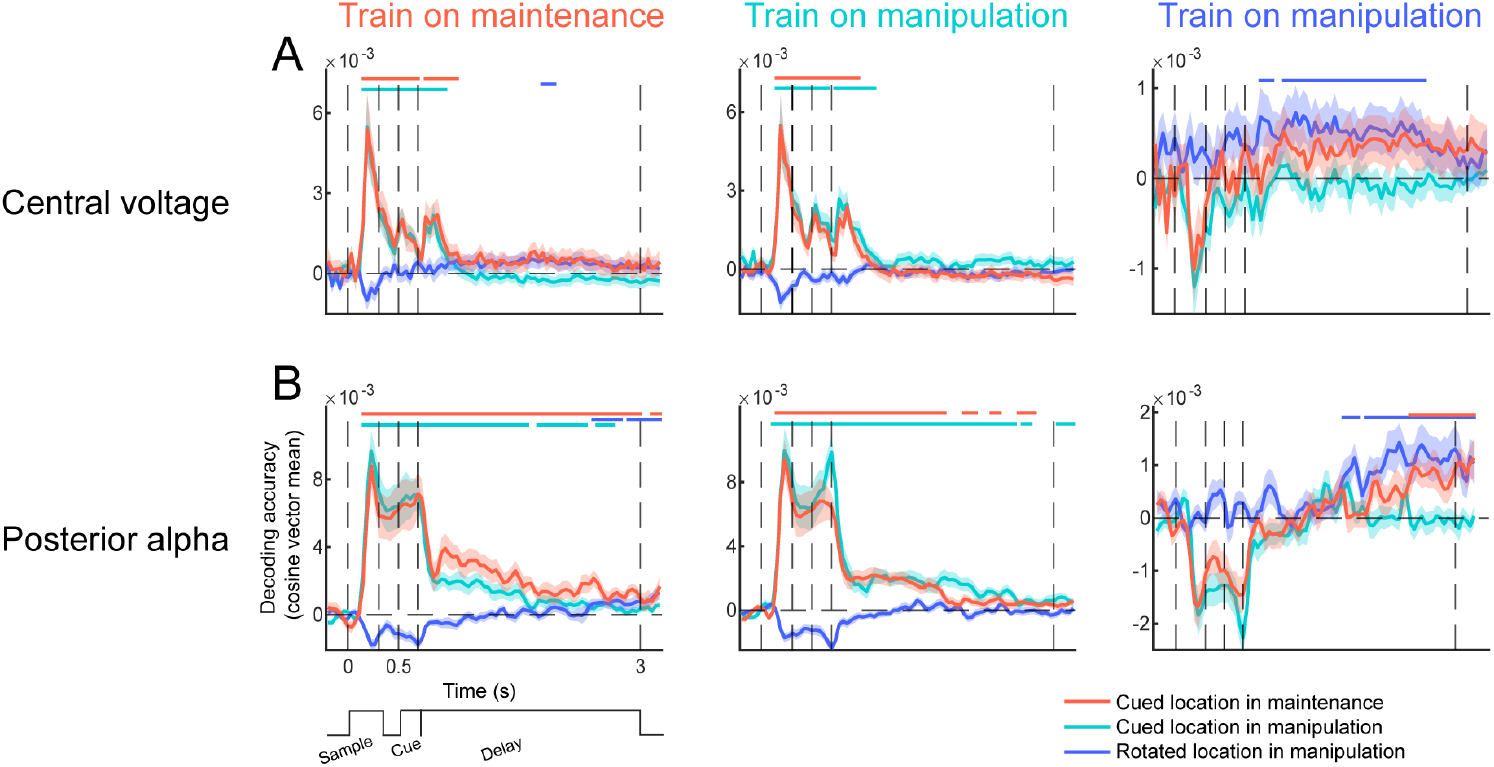
Generalization of decoding results in central voltage and posterior alpha-band (after removing influences of eye movements, N=23). (A) Generalization of location decoding between maintenance and manipulation in central voltage activity. From left to right: results from the decoder trained on the cued location in maintenance; results from the decoder trained on the cued location in manipulation; results from the decoder trained on the rotated location in manipulation. Each decoder was tested on all three conditions. Horizontal lines on top denote significance of the corresponding condition. (B). Same conventions as A, but with generalization results from posterior alpha-band activity.

**S4 Fig.**
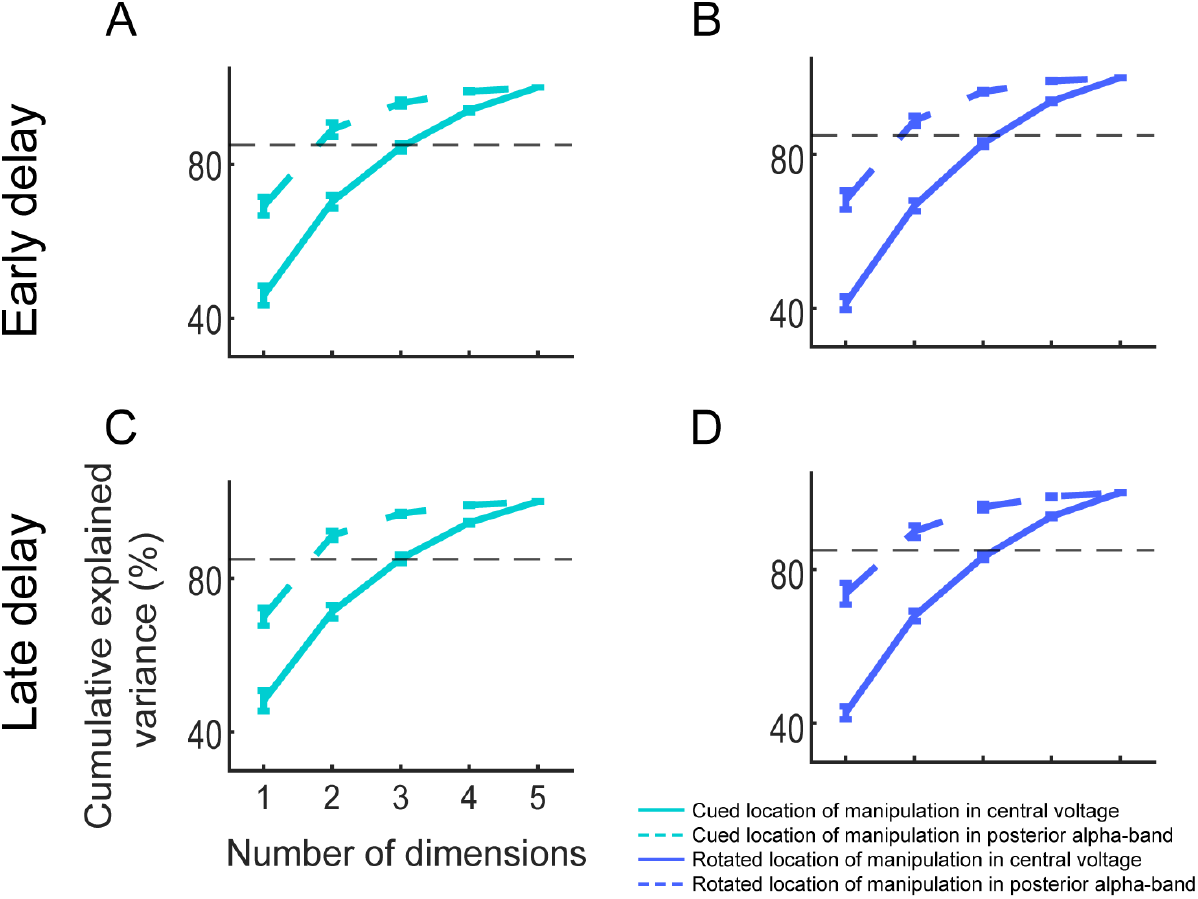
Cumulative explained variance of the subspaces (after removing influences of eye movements, N=23). (A) Cumulative explained variance as a function of number of dimensions, for the cued location in the manipulation condition during the early delay. The solid line represents results from central voltage, and the dashed line represents results from posterior alpha-band. (B) Same conventions as A, but with results from the rotated location in the manipulation condition during the early delay. (C) Same conventions as A, but with results from the cued location in the manipulation condition during the late delay. (D) Same conventions as A, but with results from the rotated location in the manipulation condition during the late delay.

**S5 Fig.**
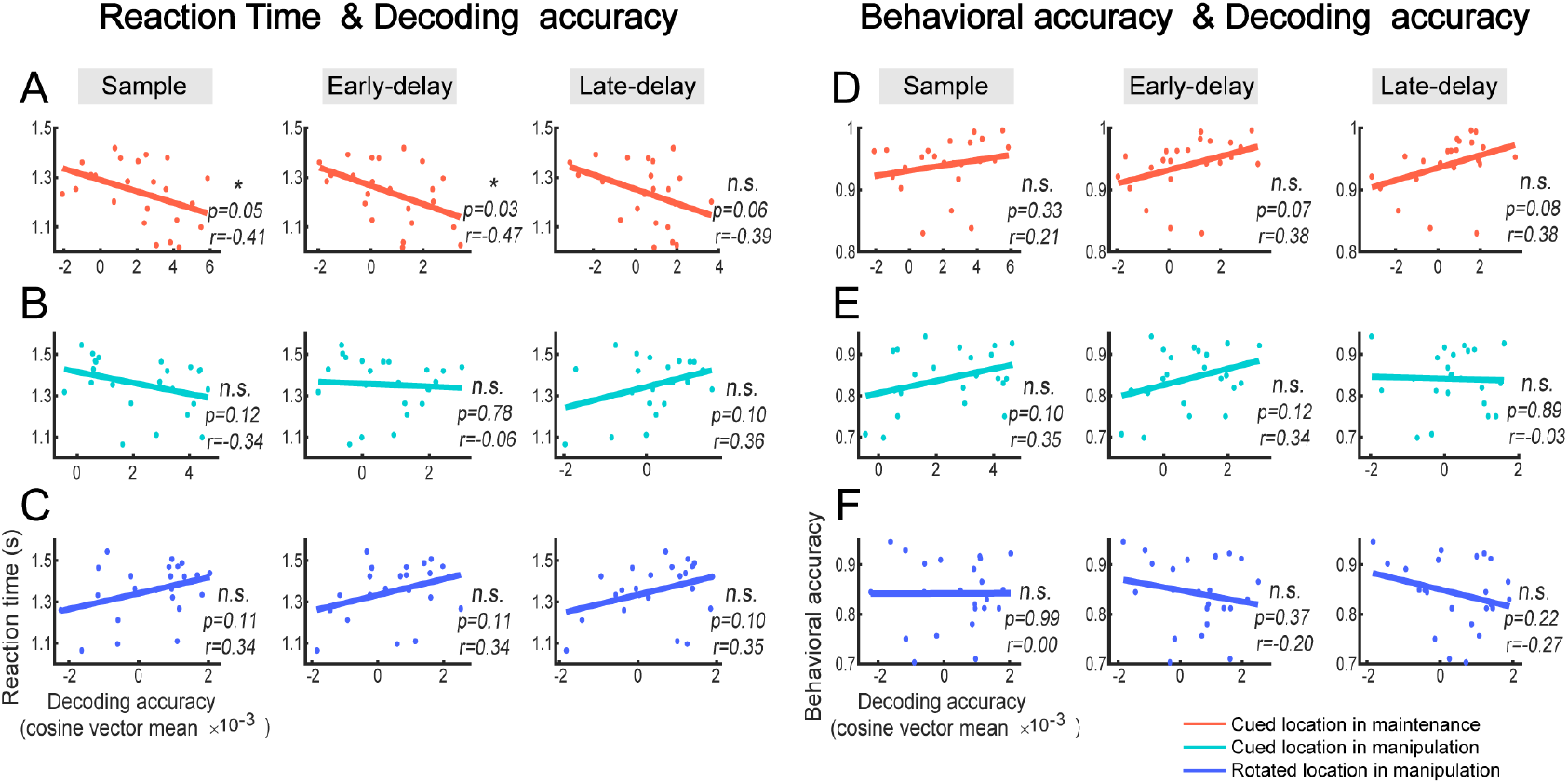
Correlation between behavioral performance and decoding results (after removing influences of eye movements, N=23). (A) Pearson correlation between reaction time and decoding accuracy of cued location in maintenance during the sample, early delay and late delay periods, respectively. (B) Correlation between reaction time and decoding accuracy of cued location in manipulation. (C) Correlation between reaction time and decoding accuracy of rotated location in manipulation. (D) Same conventions as A, but with correlation between behavioral accuracy and decoding accuracy of cued location in maintenance. (E) Correlation between behavioral accuracy and decoding accuracy of cued location in manipulation. (F) Correlation between behavioral accuracy and decoding accuracy of rotated location in manipulation.

